# High resolution spatial profiling of the hematopoietic landscape of the murine lung

**DOI:** 10.1101/2025.09.25.678618

**Authors:** Nicholas Skvir, Alexandra B. Ysasi, Veronica Jarzabek, Anthony K. Yeung, Jiaji Chen, Ruben Dries, George J. Murphy

## Abstract

Our current understanding of blood cell development and functionality stems primarily from the investigation of adult bone marrow (BM) and the fetal liver prenatally. However, emerging evidence highlights the lung as a previously underappreciated residence for hematopoietic cells. While a diversity of cells specific to the BM are known to promote the maturation and trafficking of hematopoietic cells, how the lung niche influences the development and functionality of resident cells is not known. Spatial in situ transcriptomics enables accurate mapping of cell identities and interactions within intact tissue, providing insights not accessible by dissociated single-cell profiling. Here, we present a high-resolution spatial transcriptomic atlas of the healthy adult murine lung placing specific emphasis on the hemato-endothelial landscape of this organ. As a case study, we developed a semi-automatic workflow to explicitly identify and curate rare – often multinucleated – megakaryocytes, requiring a combination of hex-binning spatial enrichment of canonical markers, expert curation, and cell boundary merging to correct for segmentation artifacts. We then characterized the spatial neighborhoods of megakaryocytes, illustrating their topological embedding within vascular, stromal, and immune microenvironments. Finally, we demonstrated the utility of this dataset for hypothesis-driven signaling studies by examining ligand–receptor interactions across pathways including BMP, VEGF, and ECM–integrin signaling. Together, this work defines the lung-blood niche and advances our understanding of the organ-specific properties of blood cells. We also provide a high-resolution spatial reference for the murine lung and demonstrate how targeted spatial in situ transcriptomics enable focused case studies of rare hematopoietic niches.

**KEY POINTS:**

1. This work represents the highest resolution gene expression mapping of the spatial symbiosis between the hematopoietic and pulmonary systems.
2. Pulmonary megakaryocytes localize within distinct vascular and stromal neighborhoods.

## INTRODUCTION

Hematopoiesis is a spatially and temporally dynamic process that has classically been ascribed to the bone marrow, which serves as the principal reservoir for hematopoietic stem and progenitor cells (HSPCs) and their downstream lineages. However, accumulating evidence demonstrates that hematopoietic activity extends well beyond the marrow and spleen, with the lung emerging as an important and previously underappreciated site of blood cell production and immune regulation. Recent discoveries have revealed that the murine lung harbors a surprisingly rich hematopoietic landscape, including resident populations of megakaryocytes (MKs), neutrophils, monocytes, dendritic cells, and innate lymphoid cells that together contribute to pulmonary immune surveillance and systemic hematopoietic function ^1–5^.

Seminal intravital imaging and transgenesis studies have demonstrated that the lung microvasculature is a site of platelet biogenesis, with significant numbers of platelets generated from MKs actively residing in and trafficking through the pulmonary circulation, particularly following insult or injury ^1,4,6^. More recently, single-cell RNA sequencing (scRNA-seq) has broadened this view by cataloging the heterogeneity of hematopoietic cells within the lung parenchyma, revealing context-dependent adaptations during infection, inflammation, and tissue repair ^7–10^. These studies underscore the concept that the lung is not only a passive filter for circulating cells but also a dynamic hematopoietic niche capable of shaping systemic immune and platelet homeostasis^11^.

Despite these advances, a critical limitation has been the ability to capture the spatial organization of hematopoietic cells within the highly complex lung microenvironment. Traditional approaches such as flow cytometry and scRNA-seq provide powerful single-cell resolution but require dissociated tissues, thereby eliminating information about tissue architecture, cellular neighborhoods, and cell–cell communication. Histologic methods can provide some spatial context but lack the multiplexing capacity required to interrogate diverse cell populations simultaneously. Consequently, our understanding of how hematopoietic cells are positioned in situ within the lung—relative to the vasculature, epithelium, and stromal compartments— remains incomplete.

Spatial transcriptomics technologies have recently transformed this landscape by enabling gene expression profiling while preserving tissue architecture^12,13^. Among these, multiplexed error-robust fluorescence in situ hybridization (MERFISH) provides the highest detection sensitivity and available resolution, allowing single-molecule detection of hundreds to thousands of transcripts in intact tissue sections^14,15^. When coupled with customized probe panels and advanced computational tools, MERFISH enables unprecedented opportunities to dissect the composition, spatial distribution, and functional interactions of rare and morphologically heterogeneous hematopoietic cells in complex tissues such as the lung.

Here, we apply MERFISH through the MERSCOPE platform to generate the highest-resolution spatial transcriptomic map of the murine lung hematopoietic niche to date. We designed a novel 500-gene probe set that captures the broad cellular diversity of the pulmonary environment, with focused enrichment for hematopoietic subsets and immunoregulatory pathways. Importantly, we developed stringent analytic pipelines to identify and characterize lung-resident MKs, a rare and morphologically complex population that plays a central role in platelet generation and local immunoregulation. By integrating high-content imaging, custom segmentation approaches, and nearest-neighbor analyses, we delineate the spatial architecture and molecular signaling networks of the lung MK niche with unprecedented resolution. Together, these data establish a foundational resource for understanding the spatial dynamics of pulmonary hematopoiesis and provide a framework for investigating how this niche is remodeled in health and disease.

## METHODS

### Tissue isolation and processing

For isolation of lung tissues, mice were euthanized via isoflurane overdose followed by transcardial perfusion to flush blood from the pulmonary system. Lungs were inflated through the trachea using 1:1 OCT:PBS (1X PBS diluted from 10X PBS using nuclease-free water). Lung lobes were bluntly dissected at the bronchi, flash frozen in OCT, and stored at -80C. Cryosectioning and slide preparation was performed in accordance with Vizgen tissue preparation guidelines. Briefly, tissue blocks were equilibrated to -20C for 30 minutes, then 10 μm sections were mounted on Vizgen MERSCOPE slides (Vizgen, Cat. 20400106) and re-frozen at -20C for 30 minutes. Slides were then incubated in pre-warmed 4% PFA fixation buffer (prepared using 32% PFA, 10X PBS and nuclease-free water) for 1 hour at 37C, washed 3X in 1X PBS (prepared using nuclease-free water), then incubated at 4C overnight in 70% ethanol to permeabilize the tissue. Autofluorescence was quenched using a MERSCOPE photo bleacher (Vizgen, Cat. 10100003) for at least 3 hours. Cell boundaries were stained using the MERSCOPE Cell Boundary Stain Kit (Vizgen, Cat. 10400118). Slides were then incubated with a custom gene panel mix at 37C for 36-48 hours according to the Vizgen Encoding Probe Hybridization Protocol. Gel embedding and clearing of the hybridized tissue was performed according to Vizgen Sample Prep Kit (Vizgen, Cat. 10400012) guidelines. Sample RNA quality was validated by extracting RNA from approximately 50 μm of sectioned tissue using an RNA 6000 Pico Kit (Agilent, Cat. 5067-1513) and analyzing with an Agilent Bioanalyzer 2100. Samples used for MERSCOPE analysis had a minimum RNA integrity number (RIN) of 8. Sample quality was further validated using the MERSCOPE Sample Verification Kit (Vizgen, Cat. 10400008) with probes targeting the mouse housekeeping gene Eef2 in a rapid imaging protocol in laser channels 561 and 647.

### Panel design

The gene panel was designed for the identification of mouse lung and airway cells with an emphasis on functional characterization of epithelial and innate immune populations. A preliminary gene panel was generated using cell-type specific markers, as well as growth factors, cytokines, chemokines, and regulatory factors associated with key signaling pathways. Highly abundant genes were excluded from the panel to avoid optical crowding, with individual gene abundances limited to <700 FPKM. Genes with fewer than 50 target regions were excluded to ensure high target specificity. The panel was further refined by excluding functionally overlapping genes and limiting the 500 gene panel to <9000 FPKM. This custom panel of genes can be found in its entirety in **Supplemental Table 1**.

### Software and computational environment

All computational analyses were performed utilizing the Giotto spatial transcriptomics software suite^16^ (v4.2.1) in RStudio Server (R version 4.4.0, RStudio version 2024.04.2-764). Analyses were run on a high-performance computing cluster environment at Boston University (BU SCC), and all parameters, intermediate objects, and scripts logged to ensure reproducibility. Additional packages loaded included gcc (v12.2.0), gdal (v3.6.4), geos (v3.11.1), hdf5 (v1.8.21), java (v11.0.4), proj (v9.2.0), and sqlite3 (v3.37.2). Scripts used to perform different stages of analysis can be found at https://github.com/nskvir/MERSCOPE-Lung-Profiling.

### Data processing and quality control

Both raw spatial outputs from Vizgen MERSCOPE, including per-transcript coordinate tables, cell segmentation boundaries, and z-stack mosaic images (DAPI, Cellbound, PolyT) were loaded into Giotto. Counts were generated by point-in-polygon overlap per z-stack (calculateOverlap/overlapToMatrix) and summed into an aggregate layer (aggregateStacks) prior to analysis, which was set as the active spat unit. Quality control filtering was performed with filterGiotto after total expression was examined for each sample (**Supplemental Figure S1**), requiring genes to be detected in at least five cells (feat_det_in_min_cells = 5), cells to contain at least five detected features (min_det_feats_per_cell = 5), and a minimum expression threshold of 1 count per feature. Normalization was performed with normalizeGiotto, applying library size normalization and scaling counts to a factor of 1,000 per cell. These thresholds were selected to balance stringency with retention of rare cell populations, given the targeted nature of the 500-gene panel.

### Dimensionality reduction, clustering and annotation

Dimensionality reduction was performed using principal component analysis (PCA), and top principal components were retained based on scree plot visualization for downstream analysis. UMAP was applied to the reduced PCA space to generate two-dimensional embeddings for visualization purposes. Unsupervised clustering was carried out with the Leiden algorithm (doLeidenCluster) using Giotto defaults, and cluster-specific marker genes were identified with the standard Giotto pipeline. Although this approach yielded multiple interpretable peripheral clusters across our samples, there remained some ambiguity among central clusters. To improve confidence in cell type annotation, we utilized a reference dataset (the LungMAP murine single-cell reference)^17^, containing approximately 40 annotated cell types, onto the MERSCOPE UMAP embeddings using newly implemented Giotto functionality. More specifically, features were intersected between the single cell and spatial data to make a joined object, which was then re-processed (filterGiotto, normalizeGiotto, runPCA and runUMAP). Lastly, Harmony integration^18^ was performed (runGiottoHarmony) followed by label transfer (labelTransfer), and re-addition of the cell metadata (addCellMetadata). This projection enabled re-labeling of cells according to their proximity to reference-defined clusters, yielding a higher-confidence annotation set that was used for all subsequent analyses.

### Megakaryocyte identification and curation

Because megakaryocytes (MKs) are both transcriptionally similar to platelets and morphologically atypical (large, often multinucleated), they posed challenges for automated identification and subsequent segmentation in the dataset. To overcome this, we applied a two-step strategy. First, we implemented a stepwise approach to identify potential regions with MKs. More specifically, we used Giotto’s tessellate function to overlay hexagonal bins across each sample, with ‘shape_size’ values set between 50–100 and calculated enrichment scores for each bin based on six canonical MK marker genes (Gp1ba, Itga2b, Mpl, Pf4, Tubb1, Vwf), using both scaled means and rescaled sums. Candidate regions were systematically screened and prioritized if both scoring approaches were positive. These regions were then examined visually by experts to discern MKs from platelet clusters, based on morphology (DAPI and cellbound stains) in tandem with robust expression and co-localization of our six marker-gene transcriptional overlays. Secondly, to correct for segmentation errors (where single MKs were often split into multiple polygons, thus identifying them as separate cells), we re-assessed cell boundaries of each putative MK. Polygons corresponding to each curated MK were merged within the Giotto object, utilizing their individual polygon IDs, new centroids were calculated, and the corrected polygons were reinserted into the Giotto object. Lastly, boundaries were redrawn using the draw function from the Terra package to ensure proper calculation of transcriptional overlaps for each cell. This ensured a one-to-one correspondence between curated MKs and polygon objects, necessary especially for downstream neighborhood enrichment analyses of rare cell types (see results).

### Proximity enrichment analysis of Megakaryocytes

Utilizing the createSpatialNetwork and cellProximityEnrichment functions within the Giotto framework, we constructed a spatial Delaunay network of all cells across a merged object comprised of both samples together in order to perform proximity enrichment analysis. Globally, pairwise enrichment scores were calculated for every cell type combination relative to a randomized null distribution. For focus on curated MKs specifically, neighborhood composition was quantified at both broad and specific levels in radial bins extending outward in 10 μm increments to determine relative cell-type proportions of neighbors at varying distances. First, a KNN network was constructed using the createSpatialKNNnetwork function. Next, by using extracted metadata, cell connections with our megakaryocytes were binned by distance, with counts and relative proportions used to construct bar plots.

### Ligand receptor analysis

To visualize potential intercellular signaling, we used Giotto’s spatCellCellcom function to calculate ligand-receptor (L/R) enrichment scores across the annotated lung cell types. A curated set of L/R pairs was derived from our custom 500-gene panel, which included genes spanning angiogenesis, BMP signaling, chemokine/chemotaxis, apoptosis, extracellular matrix, growth factors, and interferon pathways. The analysis was run with 1,000 permutations to generate null distributions for comparison. Enrichment was quantified as the fold change between observed versus randomized ligand–receptor co-localization, with significance assessed by empirical *p*-values from the permutations. To generate our dot plots summarizing enrichment across L/R pairs, representative cell pairs showing elevated log fold change across 11 selected L/R pairs from our custom panel were taken from the top 60 entries in the interactions table generated from the previous step, sorted by adjusted p-value.

## RESULTS

### High resolution overview of the hematopoietic, endothelial and pulmonary landscapes of the adult murine lung

To establish a high-resolution reference map of the healthy murine lung, we generated the highest possible spatial transcriptomic resolution map of intact lung tissues by using multiplexed error-robust fluorescence in situ hybridization (MERFISH)^14,19^ through the MERSCOPE platform. To accomplish this, we used previously described scRNA-seq datasets^20–22^ to guide the development of a novel 500 gene MERFISH probe set designed to maximally cover the complex lung niche composition and inherent biological variability. This included markers for different cell types and subtypes (i.e. pulmonary, hematopoietic/immune, endothelial, etc.) and further enrichment for genes involved in immunoregulatory processes, ligand-receptor signaling, and developmental pathways. We profiled two independent tissue blocks with this approach; in total, 184,911 cells were captured across both samples (67,872 in Sample A and 117,039 in Sample B). Following quality control and reference-based annotation, the dataset resolved into 38 discrete cell types spanning all major compartments of the adult lung (**Fig. 1A**). These included epithelial populations such as alveolar type 1 & 2 cells (AT1/AT2), ciliated, and Sox9+ epithelial cells; endothelial subsets including CAP1/EPC, CAP2 (‘aerocyte’ capillary), arterial, venous, and lymphatic endothelial cells (AEC, VEC, LEC); mesenchymal lineages including alveolar fibroblasts (AF1, AF2), secondary crest myofibroblasts (SCMF), and mesothelial cells; and a diverse array of immune and hematopoietic populations, including macrophages (alveolar and interstitial), dendritic cells (maDC), lymphocytes (ILC, T-cells, B-cells, NK cells, Tregs), erythroid cells, and megakaryocyte/platelets. This global view revealed detailed architecture of the adult lung, including conducting airways, alveolar regions, and interstitial niches, as well as clearly resolved mesothelium and lymphatic vasculature. Within conducting airways, ciliated epithelial cells were clearly resolved, surrounding the airway lumen (**Fig. 1B**). In the adjacent interstitium, immune cell aggregates could be detected, illustrating the ability of MERSCOPE to localize hematopoietic populations with high spatial fidelity (**Fig. 1B, red arrow**). Together, these results demonstrate that MERSCOPE, combined with a targeted transcript panel, can resolve both broad tissue architecture and rare hematopoietic niches *in situ*. This dataset thus serves as a valuable reference framework for interrogating murine lung biology in situ at the highest level of resolution.

**Figure 1.**
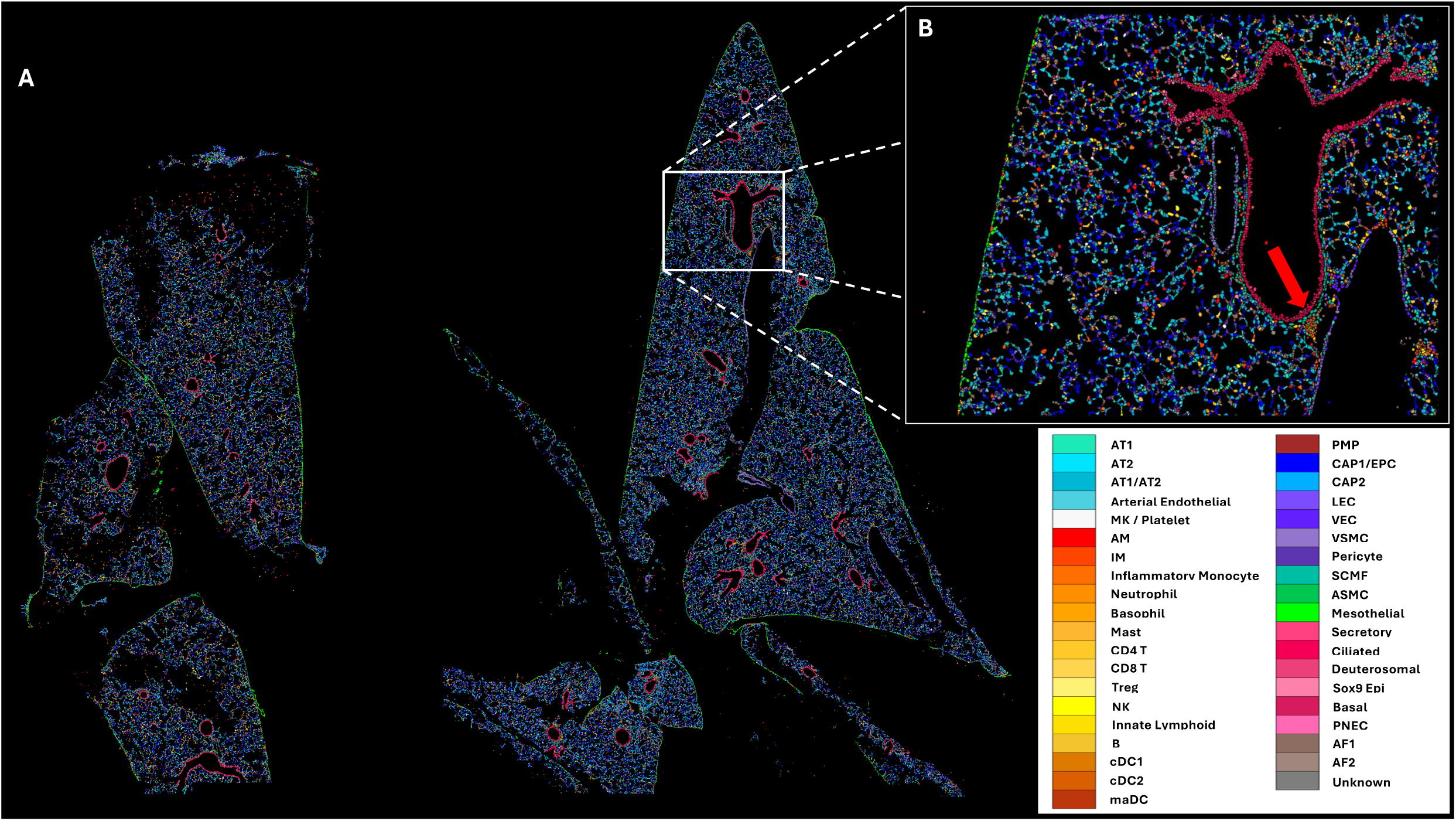
High resolution overview of the hematopoietic, endothelial and pulmonary landscapes of the adult murine lung. A) The combined spatial overview of two distinct murine lung samples that were profiled using a unique 500 gene panel combining B) Enlargement highlighting ciliated cells surrounding a large conducting airway. The red arrow denotes a cluster of immune cells in the interstitium localized to the lower border of the conducting airway.

### Single cell deconstruction of MERFISH spatial data identifying 38 individual cell types

To classify the transcriptionally distinct populations within our dataset, we first applied standard Giotto^16^ workflows. PCA followed by UMAP embedding and Leiden clustering yielded ∼17-19 clusters across both samples. While many peripheral clusters could be annotated based on canonical marker expression, a substantial fraction of cells fell into a large, ambiguous central cluster, suggesting that Leiden clustering alone was insufficient to resolve the full diversity of cell types seen in the lung (**Supplementary Fig. S2**). To improve confidence in cell annotations, we leveraged the LungMAP murine single-cell reference^17^ (CellRef), which contains approximately 40 well-curated cell types from healthy mouse lung. We used Harmony^18^ integration to create a shared embedding space between the single-cell reference and spatial data followed by label transfer of annotated CellRef data to cell identities in the spatial data (**Fig. 2A**). Using this refined approach, the MERFISH dataset resolved into nearly 40 discrete cell types spanning epithelial (AT1, AT2, ciliated, Sox9+), endothelial (CAP1/EPC, CAP2, arterial, venous), fibroblast/mesenchymal (AF1, AF2, SCMF, mesothelial), immune (myeloid, lymphoid), and hematopoietic (MK/platelet, erythroid) lineages (**Fig. 2B**). Altogether, this CellRef-informed approach substantially improved the granularity of cell annotations compared to unsupervised clustering and was used as the foundation for all downstream analyses.

**Figure 2.**
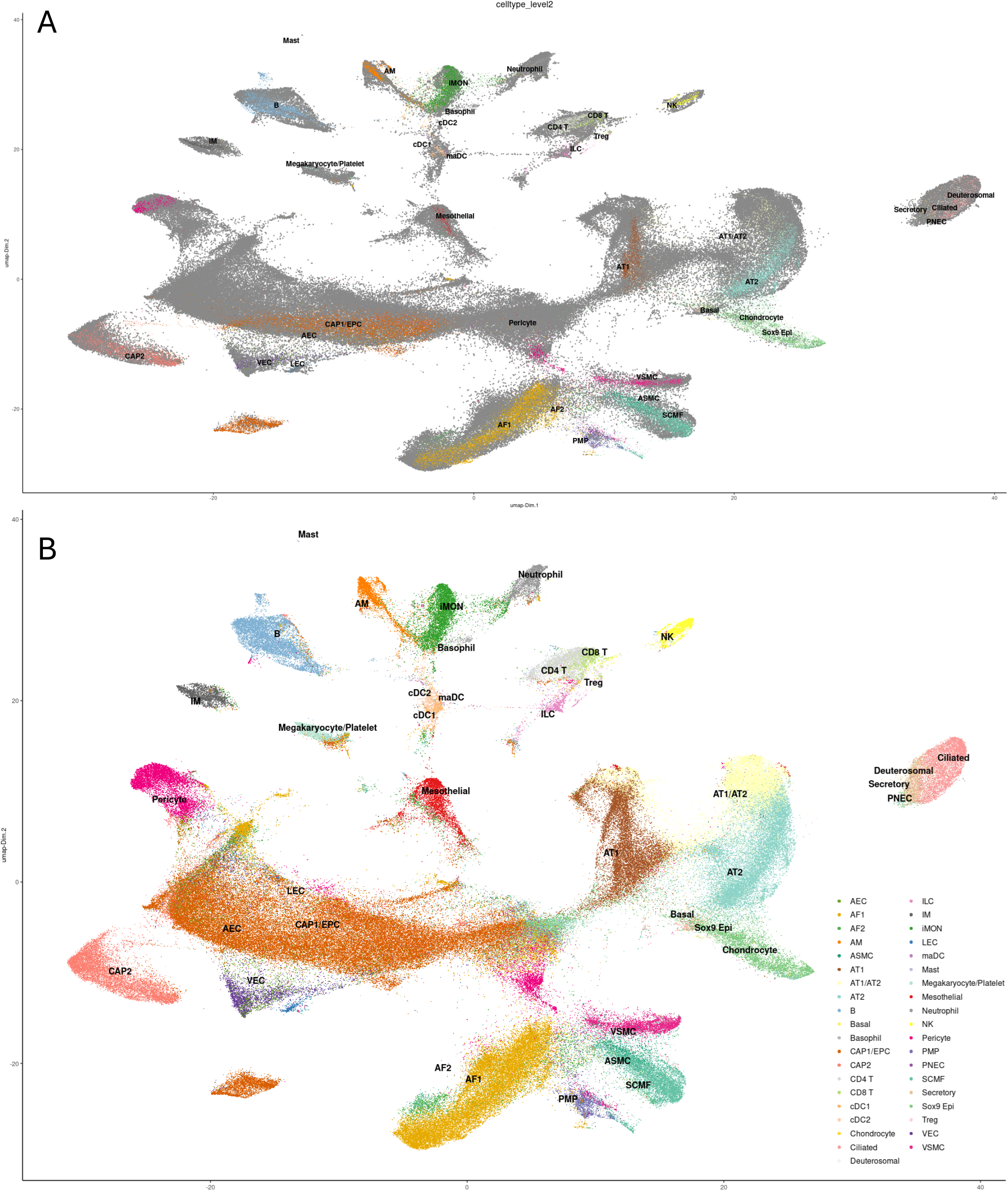
Merged-sample UMAPs and scRNA-based application of cell labels. A) This figure shows the scRNA CellRef data (colored) projected on top of our MERSCOPE data (grey) and how they align. B) The UMAP after subsequent label application onto our samples, improving the resolution of our calls and allowing for the identification of nearly 40 discrete cell types across our data sets.

### High resolution spatial profiling of the lung and megakaryocyte niche

Megakaryocytes (MKs) are rare but biologically significant hematopoietic cells previously reported to reside in the lung^4,23,24^. Because of their scarcity, large size, and transcriptional similarity to platelets, MKs could not be cleanly resolved using the initial default clustering approach. Hence, we developed a more explicit and semi-automatic pipeline to identify MKs with high confidence. First, the spatial transcriptomics data was rasterized using a ‘hex-binning’ approach across our two datasets (detailed in the methods section) with the goal to identify tiled regions showing enrichment of MKs based on, six canonical MK marker genes (Gp1bα, Itga2b, Mpl, Pf4, Tubb1, Vwf). These enriched regions were then prioritized as potential candidate MK locations (**Fig. 3A**) and further confirmed in the following manner. First, DAPI and cellbound staining was used to identify characteristic MK morphology, including large, multinucleated cells (**Fig. 3B–C**). Then, transcript overlays were used to visualize marker co-expression and provide final confirmation of MK identity. To correct for initial segmentation errors, due to large – and often multinucleated – single MKs being split into multiple polygons, we manually corrected these cell boundaries where necessary, and used these corrected cell polygons for downstream analysis in the Giotto analysis framework (**Fig. 3D**). This workflow enabled us to conservatively identify genuine MKs across both samples, providing a foundation for downstream neighborhood analyses of the megakaryocyte niche within the lung, while also underscoring the potential complexity in analysis of cell types with unique morphologies. Thus, we next examined the spatial context of MKs within the lung. Using our reference-based cell type labeling, we constructed a spatial network of all cells across both samples and performed proximity enrichment analysis. Firstly, pairwise enrichment plots (Fig. 4A) were generated for the whole lung, as well as corresponding heatmaps and cell proximity networks (**Supplementary Fig. S3**). Preliminary global neighborhood enrichment of the top 25 enriched and bottom 25 depleted cell type pairings identified both expected and novel relationships between lung cell types. The strongest enrichments were generally homotypic, with cells preferentially adjacent to others of the same type (e.g., deuterosomal–deuterosomal, etc). These patterns likely reflect conserved tissue organization and lineage clustering. Additional enriched proximities included ciliated– deuterosomal pairs and cDC1–cDC2, consistent with developmental and functional links among these cell types, as well as structural associations such as airway smooth muscle–secondary crest myofibroblast (ASMC–SCMF) and AF2–mesothelial. In contrast, the most depleted interactions often involved secretory or ciliated epithelial cells paired with unrelated lineages (e.g., lymphatic endothelial, NK cells, AT1, etc.), underscoring the compartmentalization of airway-associated epithelia away from vascular, immune, and hematopoietic niches. Together, these findings reinforce expected features of lung organization while also highlighting less well-characterized associations. For example, enrichment of AF and mesothelial cells reflects a known connection, specifically in lung injury and disease^25,26^ while associations between cell types such as ASMC–SCMF suggests structural or lineage-linked interactions not typically described in adult murine lung. Similarly, proximity of dendritic cells to innate lymphoid cells and interstitial macrophages supports immune–immune cross-talk within interstitial niches. While some of these interactions align with known developmental or functional relationships (e.g., ciliated–deuterosomal, cDC1–cDC2), others may represent underappreciated aspects of stromal and immune organization revealed by spatial profiling.

**Figure 3.**
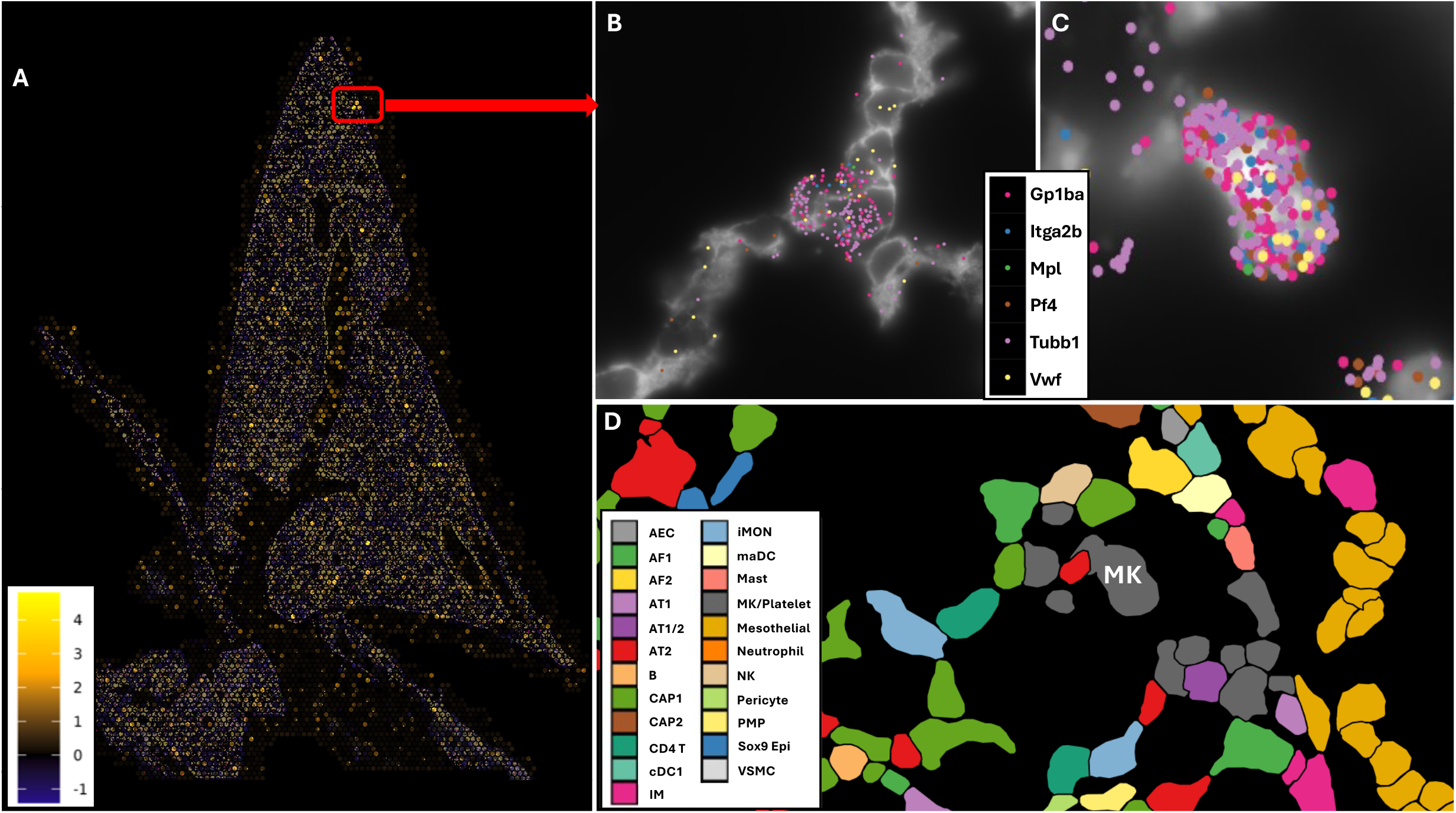
Transcription-based identification of putative megakaryocytes within the murine lung. A) To locate specific instances of megakaryocytes, a hex-binning approach was used, whereby the whole image was subset into hexagonal compartments. The color of each compartment was determined by the relative enrichment of six marker genes (depicted in B & C). Compartments with the brightest colors were zoomed in on (one such cluster of compartments indicated by the red box), and corresponding regions were assessed for the presence of megakaryocytes. Coloration was performed based both on scaled means (pictured here) as well as rescaled sums (**Supplemental Fig. S4**). B & C) Representative zoomed images (A: Cellbound, B: DAPI) of megakaryocytes located from A. Colored dots denote six individual transcripts used as markers– Gp1ba, Itga2b, Mpl, Pf4, Tubb1 and Vwf. D) Polygons were drawn for each cell boundary, and colored by cell type as assigned by label transfer based on an external, gold standard single-cell RNA sequencing dataset. These calls were used for subsequent statistical analysis.

**Figure 4.**
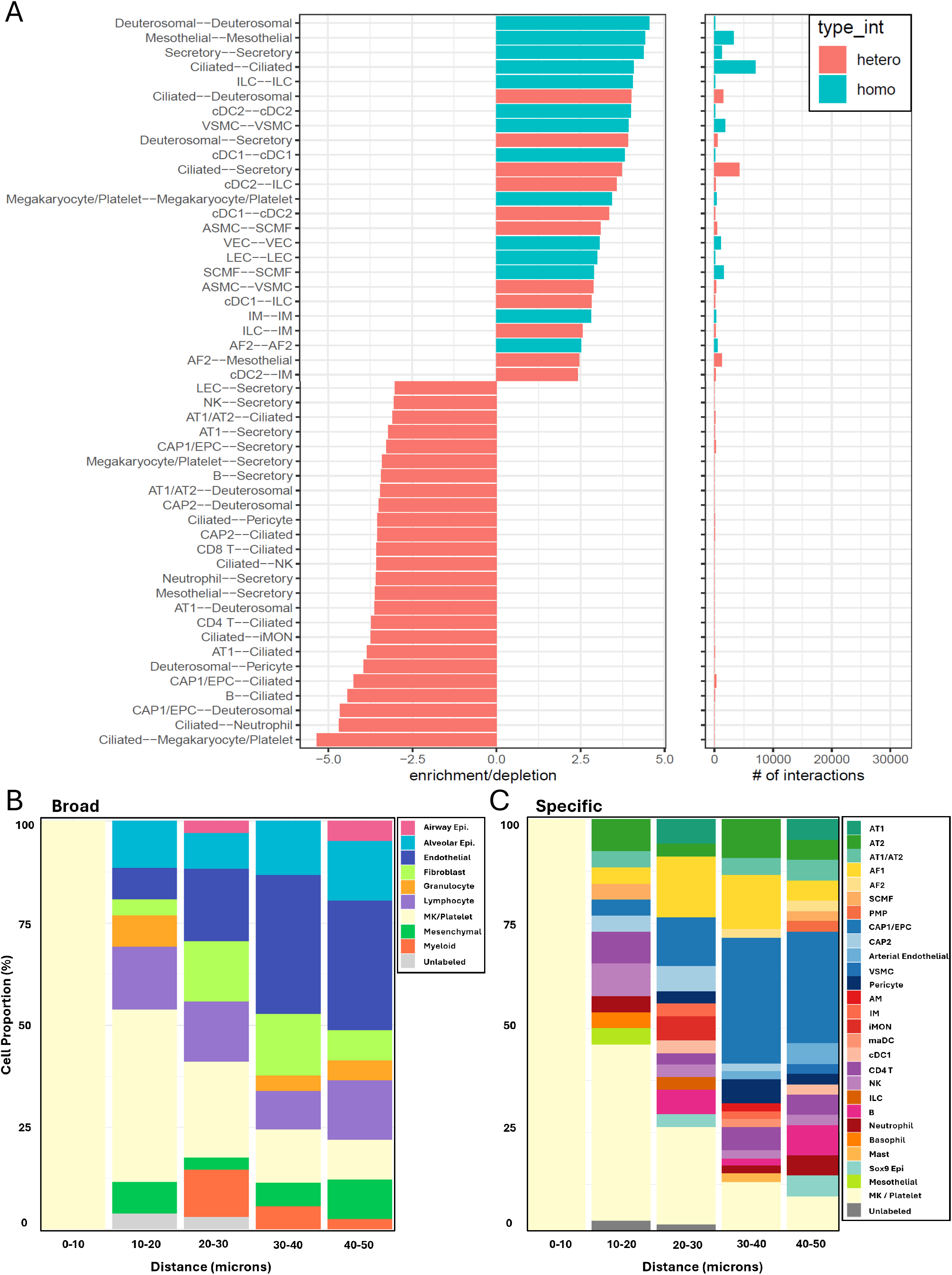
Global and megakaryocyte-specific neighborhood enrichment in the murine lung. A) Utilizing our scRNA-based labeling, cells in close proximity were profiled for their relative enrichment or depletion relative to a null distribution to observe potential novel findings pertaining to the immune landscape of the murine lung. Each cell type was compared in a pairwise fashion to every other cell type within the analysis, and bar plots were drawn showing the top 25 most enriched and depleted cell pairings within our two samples, based on cell-cell proximity scores. Blue bars show enrichment/depletion between same cell types, while red bars show the same between differing cell types. B-C) Distance-binned neighborhood composition surrounding curated megakaryocytes, showing immediate MK/platelet neighbors and enrichment of endothelial, alveolar epithelial, and immune cell types within 20–50 μm. The two plots reflect both broad (B) and more specific (C) cell typing based on our reference labels from LungMAP.

Next, to examine the local niches of our validated MKs specifically, we looked at the subset of cell proximity interactions in which they were involved. From this list of interactions, we generated bar plots showing proportions of cells in close proximity to MKs, binned by radial distance (**Fig. 4B,C**). We did this at two levels, using both broad and more specific cell identities from our earlier reference-based labeling. While at the closest distances (0–10 μm), MKs were almost exclusively neighbored by other MK/platelets, at 10–20 μm, a mosaic of additional cell types emerged, including alveolar epithelium (AT1/AT2), capillary endothelial cells (CAP1/EPC, CAP2), and lymphocytes, with smaller contributions from fibroblasts, granulocytes, and mesenchymal populations. Beyond 20 μm, endothelial representation increased further, consistent with vascular association, while airway epithelium and myeloid cells appeared at low-to-moderate levels. CAP1/EPC progressively becomes the largest non-MK/platelet component, while AF1 remains a steady, medium-sized fraction; additional minor slivers (e.g., AM, cDC1, Sox9+ epithelium) appear without overtly displacing other types. Notably, AT1/AT2 together remain represented at all distances, indicating that MKs are consistently positioned within alveolar parenchyma—near, but not exclusively abutting, the epithelial surface. Taken together, these observations motivate a model in which pulmonary MKs are supported by overlapping endothelial, epithelial, stromal, and immune cues. Specifically, the persistent alveolar epithelial contribution suggests epithelial-derived factors may regulate MK maturation or platelet release, while endothelial enrichment is compatible with vascular positioning and potential platelet egress. Stromal and immune neighbors may provide additional trophic or regulatory inputs.

### Examination of ligand-receptor interactions within the lung

To further examine modes of cellular communication within this dataset, we interrogated ligand– receptor (L/R) signaling pathways represented in the custom panel. Using Giotto’s spatCellCellcom functionality, we generated a table of putative L/R interactions across all annotated cell types (**Supplemental Table S2**). From this, we visualized a representative set of interactions in a dot plot (**Fig. 5A**). Dot size corresponds to empirical significance, while color indicates relative fold-change enrichment compared to randomized null distributions. This analysis recapitulated several well-established signaling axes, including angiogenesis (Vegfa– Flt1/Kdr), chemokine signaling (Ccl2–Ccr2), BMP pathways (Bmp4–Bmpr2), matrix–integrin interactions (Col1a1–Itga1), apoptosis (Fas–FasL), and interferon response (Ifna–Ifnar1).

**Figure 5.**
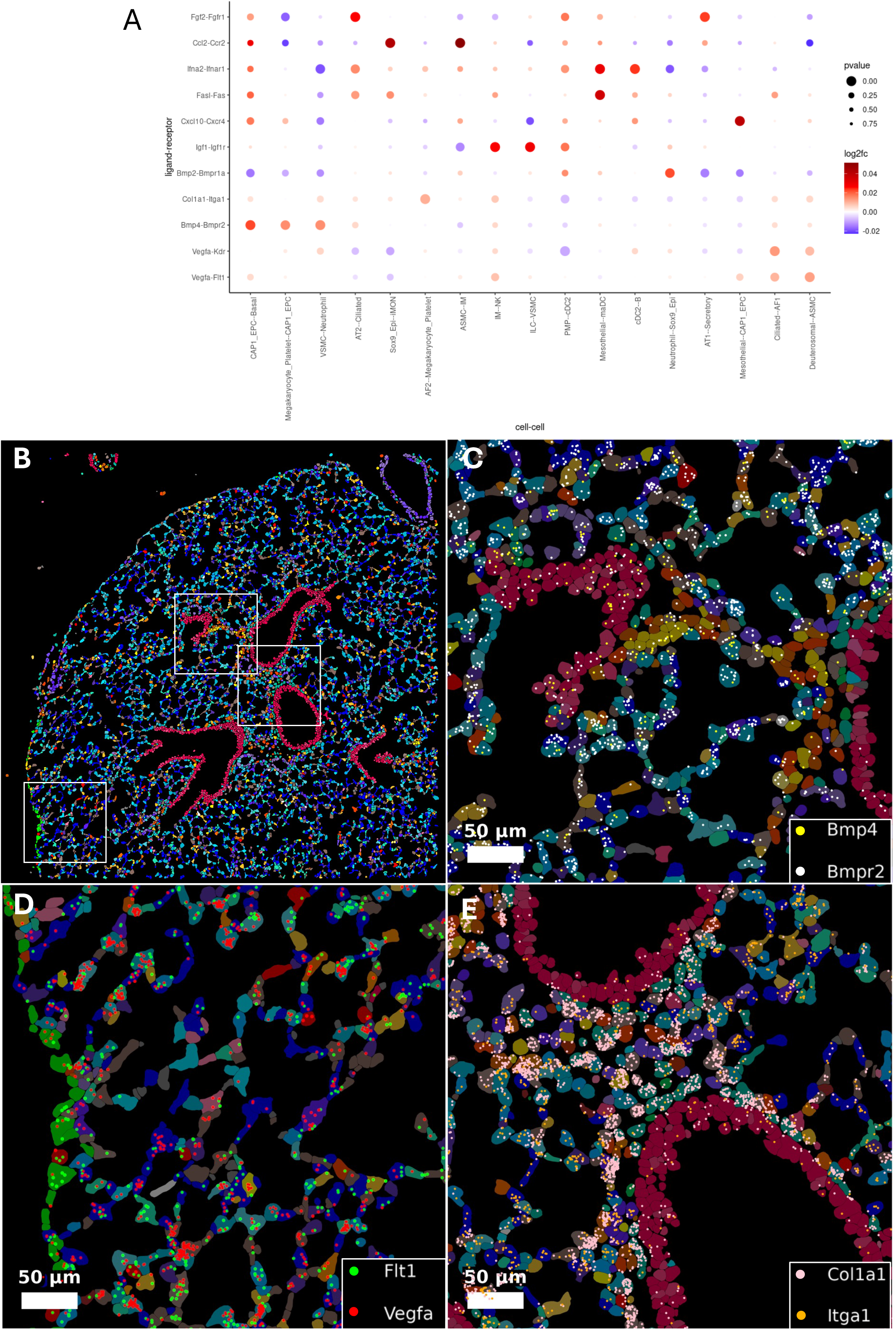
Ligand/Receptor enrichment and depletion between cell types. A) A dot plot showing relative enrichment and depletion of 11 representative types of ligand/receptor signaling between cell types within our samples, derived from a table of spatial cell-cell communication scores. The size of the dots reflects the significance, while the color represents relative fold change of observed interactions compared to a hypothetical null distribution over 1000 iterations. B) An overview of a subsection of tissue from sample B, displaying representative zoomed views of several types of L/R signaling, showcasing the capabilities of MERSCOPE to resolve each respective transcript at sub-cellular resolution. **(C) Bmp4–Bmpr2 (BMP signaling): Bmpr2 receptors (white)** were abundant in CAP2 endothelial, AT1/AT2 epithelial, pericytes, and venous endothelial cells, while **Bmp4 ligands (yellow)** were less frequent but present in AT1/AT2, AF2 fibroblasts, ILCs, and ciliated cells. Occasional junctions, such as AF2–epithelial boundaries, demonstrated potential stromal-to-epithelial signaling. **(D) Vegfa–Flt1 (angiogenesis): Vegfa ligands (red)** were observed around ciliated/deuterosomal cells and within AT1/AT2 epithelium, fibroblasts (AF1/AF2), mesothelial cells, and CAP1/EPC endothelium. **Flt1 receptors (green)** were most prominent in AT1/AT2 and AF2 fibroblasts, with additional expression in mesothelial and endothelial subsets. Ligand and receptor co-localization in AT1/AT2 and AF2 suggests autocrine signaling, while complementary expression at epithelial–endothelial junctions supports paracrine communication. **(E) Col1a1–Itga1 (matrix–integrin interactions): Col1a1 ligands (pink)** were abundant in AF2 fibroblasts, SCMFs, and immune subsets (ILCs, cDC1, cDC2), whereas **Itga1 receptors (orange)** localized mainly to pericytes, VSMCs, and AT1/AT2 epithelium. Overlaps occurred at fibroblast–immune and epithelial–pericyte interfaces, suggesting collagen-mediated matrix–integrin signaling across stromal–epithelial and stromal–vascular boundaries. Together, these images highlight spatially resolved ligand–receptor cross-talk across stromal, epithelial, endothelial, and immune compartments of the lung.

To validate these statistical enrichments *in situ*, we visualized selected ligand–receptor pairs directly within the MERSCOPE images (**Fig. 5B–E**). **Bmp4–Bmpr2** signaling showed broad receptor expression across epithelial, endothelial, and mural (pericyte, VSMC) compartments, with sparse stromal and epithelial ligand distribution suggesting locally constrained activation. **Vegfa–Flt1** interactions involved both autocrine and paracrine configurations, with ligands and receptors co-localized in alveolar epithelium and fibroblasts, and complementary expression at epithelial–endothelial interfaces. **Col1a1–Itga1** highlighted stromal fibroblasts and immune cells as key collagen sources, with receptors enriched in epithelial and vascular mural cells, consistent with matrix–integrin signaling across stromal–epithelial and stromal–vascular boundaries. Together, these examples underscore the ability of spatial transcriptomics to resolve cross-compartment communication in the lung with both statistical and spatial fidelity. While not intended as a comprehensive atlas of lung signaling, these analyses demonstrate how this dataset can be mined to interrogate diverse pathways of interest.

## DISCUSSION

In this study, we present a high-resolution spatial transcriptomic map of the adult murine lung, generated using Vizgen MERSCOPE with a custom 500-gene panel. By combining two independent tissue blocks, we assembled a dataset of ∼185,000 cells spanning epithelial, endothelial, mesenchymal, immune, and hematopoietic compartments. Using reference-based annotation, nearly 40 discrete cell types were confidently identified. Our findings extend prior work that increasingly recognizes the lung as more than a passive site of gas exchange, highlighting instead its role as an active hematopoietic niche with contributions to platelet biogenesis and immune regulation. Previous studies have shown that megakaryocytes (MKs) and their progenitors can traffic to the lung vasculature, where they contribute to circulating platelet pools and engage with endothelial and immune neighbors. By adding spatial resolution, our study refines this view, identifying specialized microenvironments in the alveolar parenchyma where MKs reside. These neighborhoods were layered: MKs were embedded within a background of alveolar epithelium, neighbored closely by capillary endothelium, and consistently accompanied by fibroblast/mesenchymal populations and low-level immune surveillance. This organization underscores that pulmonary MKs are supported by overlapping epithelial, vascular, stromal, and immune cues—echoing but also extending niche dynamics described in the bone marrow.

Beyond individual findings, the dataset itself represents a community resource. Prior single-cell RNA-seq atlases, such as LungMAP, have provided invaluable reference frameworks for murine lung biology, but lack spatial information. By adding high-resolution transcript localization, this MERSCOPE dataset complements those resources and enables researchers to interrogate cell types, neighborhoods, and signaling pathways directly within their tissue context. We envision this dataset as a high-fidelity spatial map that others can mine to address diverse questions about lung organization and function. Clinically, this spatial framework provides context for understanding how pulmonary vascular injury, inflammation, or fibrosis might perturb hematopoietic dynamics. For example, disruption of endothelial or stromal support could alter platelet production, while immune remodeling of the interstitium could affect MK functionality and platelet release. More broadly, our dataset establishes a foundation for probing ligand– receptor networks at the lung–blood interface, opening avenues to identify therapeutic pathways that could be targeted to modulate thrombosis, immunity, or repair in cardiopulmonary disease. At the methodological level, this work illustrates strategies that can be generalized to other spatial studies. Hex-binning proved an effective strategy for locating rare cells when clustering alone was insufficient; projection of an external single-cell reference (LungMAP) substantially improved cell type annotations; and ligand–receptor analysis demonstrated how targeted panels can yield curated pathway-level insights, validated directly in situ. Together, these approaches highlight how spatial transcriptomics bridges histological context with transcript-level detail, enabling both robust reference mapping and focused biological case studies.

Despite these advances, our study also has limitations. The use of a targeted 500-gene panel, while providing the highest detection sensitivity and designed for breadth across lung and hematopoietic biology, necessarily restricts the scope of detectable signals relative to unbiased transcriptome-wide approaches. The small number of megakaryocytes identified limits statistical power for neighborhood comparisons, and our reliance on manual validation introduces subjectivity, though consensus review helped mitigate this. Finally, the dataset represents healthy adult murine lung tissue and does not address variation across developmental or disease states.

In conclusion, we provide both a high-resolution spatial reference of the murine lung and a case study of megakaryocyte niche analysis. By demonstrating how curated workflows can resolve rare cell types and uncover spatially organized interactions, this work underscores the potential of MERSCOPE and related technologies to advance our understanding of tissue organization.

## ACKNOWLEDGEMENTS

We acknowledge members of the Center for Regenerative Medicine (CReM) and the section of Hematology-Oncology at the Boston University School of Medicine for critical feedback on this manuscript.

## AUTHORSHIP CONTRIBUTIONS

N.S. performed experimental design, data collection, data analysis, interpretation of data and manuscript preparation. A.B.Y. performed all sample preparation, as well as design of the custom panel. She also provided expert guidance pertaining to our MK calls together with A.K.Y. G.J.M. performed and supervised experimental design, interpretation of data and manuscript preparation. V.J, J.C., and R.D. assisted with insight into troubleshooting bioinformatic and computational analyses where needed, with expertise in actively developing the Giotto platform.

## DISCLOSURES

The authors of this manuscript have no conflict of interest to disclose.

## Notes

### Competing Interest Statement

The authors have declared no competing interest.

