## Supplemental Figures for "High resolution spatial profiling of the hematopoietic landscape of the murine lung"

**Description for Supplemental Table S1**:
This table contains a curated list of all candidates for our custom 500-gene MERSCOPE panel, with corresponding gene symbol, full name/protein, and associated cell population/keywords. Each of the candidate genes were included in the custom panel unless excluded, as indicated by colored cells. Corresponding color-coded explanations for exclusion are in the second tab, ‘Exclusions’, and include categories such as ‘Untargetable’ (red), ‘Low detection efficiency’ (yellow), ‘Exceeds FPKM limit per gene (>1000)’ (purple), ‘High FPKM’ (lavender), and ‘Low priority/repetitive’ (orange).

**Description for Supplemental Table S2**:
This table is the Interactions table (sorted by p.adj values) output by the Giotto function spatCellCellcom, which gives a list of spatial cell-cell communication scores based on spatial expression of interacting cells. This table was used to generate the Fig.5A dotplot, utilizing the significance and logfc values. Explanations for the contents of each column, as well as the spatCellCellcom function itself can be found in the Giotto documentation here: https://giottosuite.com/reference/spatCellCellcom.html

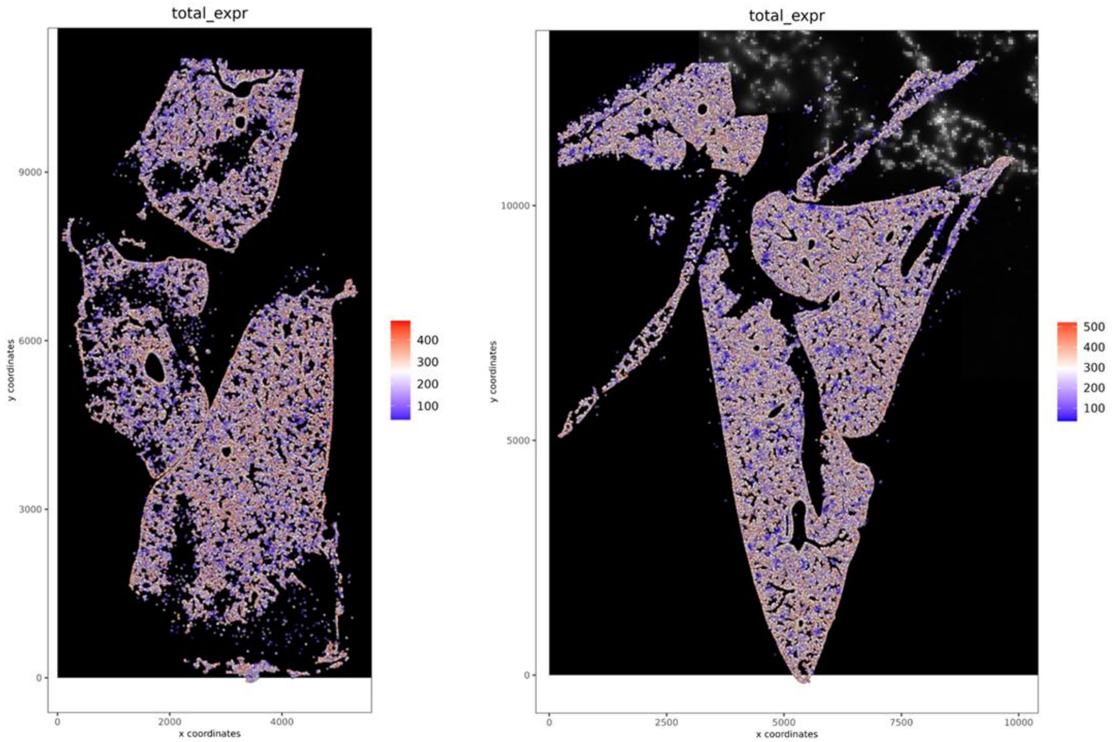


**Supplemental Figure S1.**

**Total expression distributions for each sample.** Plots for sample A (left) and B (right) showing the total number of transcripts detected per cell after aggregation. These plots provide a QC overview of capture efficiency and support downstream normalization, illustrating that both samples yielded comparable transcript detection profiles.

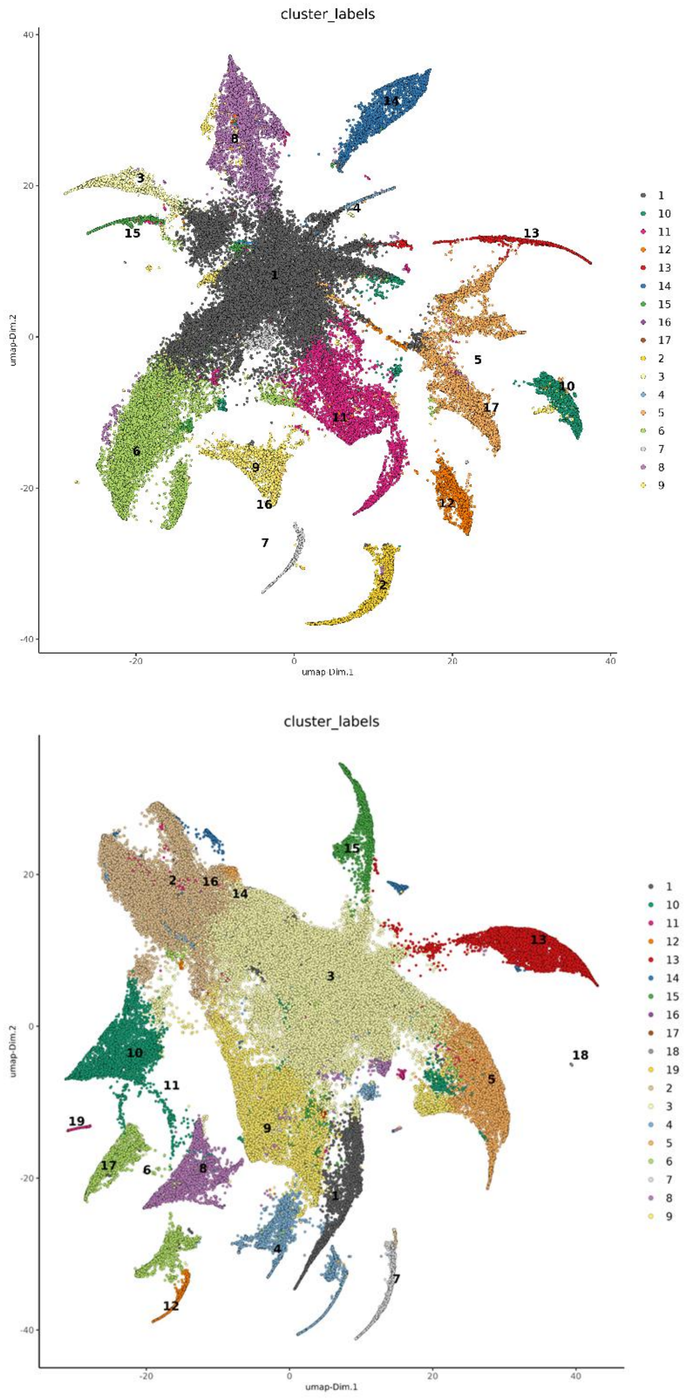


**Supplemental Figure S2.**

**Preliminary UMAPs for our two samples**. Sample A (top) and B (bottom) using built-in Leiden clustering. Clusters generated in this way contained large, ambiguous, central clusters that were not easily resolved. This prompted us to utilize an alternative approach, utilizing a reference dataset to validate our calls.

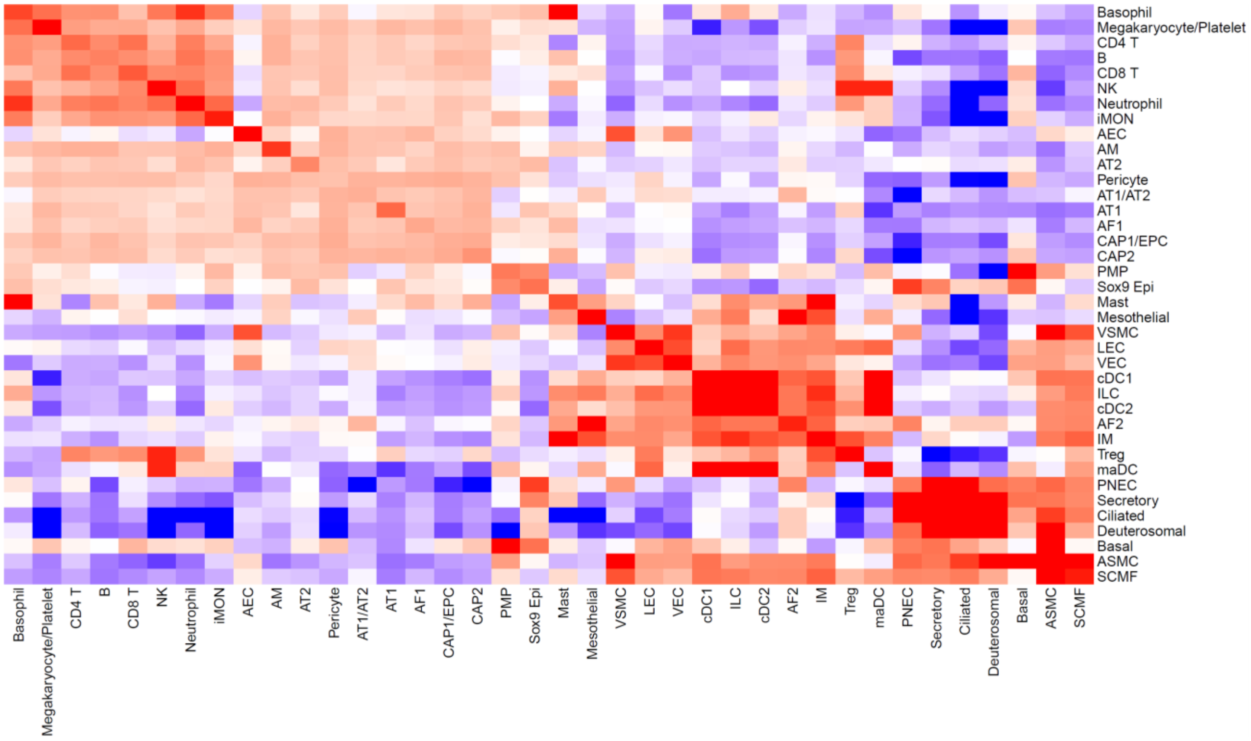
Basophil
Megakaryocyte/Platelet
CD4 T
B
CD8 T
NK
Neutrophil
iMON
AEC
AM
AT2
Pericyte
AT1/AT2
AT1
AF1
CAP1/EPC
CAP2
PMP
Sox9 Epi
Mast
Mesothelial
VSMC
LEC
VEC
cDC1
ILC
cDC2
AF2
IM
Treg
maDC
PNEC
Secretory
Ciliated
Deuterosomal
Basal
ASMC
SCMF

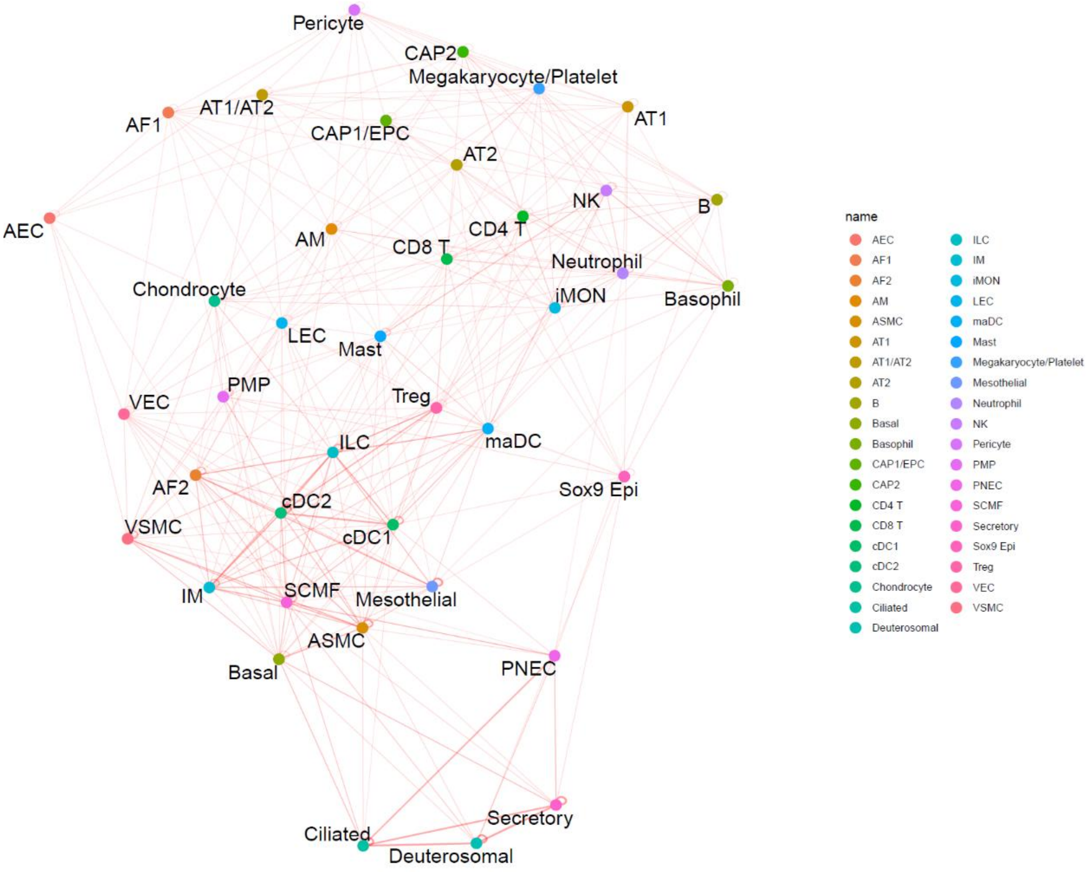
Pericyte
CAP2
Megakaryocyte/Platelet
AT1/AT2
AF1
AT1
CAP1/EPC
AT2
AEC
NK
B
CD4 T
AM
CD8 T
Neutrophil
Chondrocyte
iMON
Basophil
LEC
Mast
PMP
VEC
Treg
ILC
maDC
AF2
Sox9 Epi
cDC2
VSMC
cDC1
IM
SCMF
Mesothelial
ASMC
Basal
PNEC
Secretory
Ciliated
Deuterosomal
name
AEC
AF1
AF2
AM
ASMC
AT1
AT1/AT2
AT2
B
Basal
Basophil
CAP1/EPC
CAP2
CD4 T
CD8 T
cDC1
cDC2
Chondrocyte
Ciliated
Deuterosomal
ILC
IM
iMON
LEC
maDC
Mast
Megakaryocyte/Platelet
Mesothelial
Neutrophil
NK
Pericyte
PMP
PNEC
SCMF
Secretory
Sox9 Epi
Treg
VEC
VSMC

**Supplemental Figure S3.**

**Alternate visualizations of pairwise cell-cell enrichment and depletion.** Heatmap (top) and proximity network (bottom) visualizations of the relative enrichment or depletion of each cell type. The heatmap shows each pairwise combination cells from among the nearly 40 identified cell types with our CellRef-based labeling methodology, while the proximity network shows similar relationships with line weights corresponding specifically to enrichment (with NA & Unknown labels excluded from each).

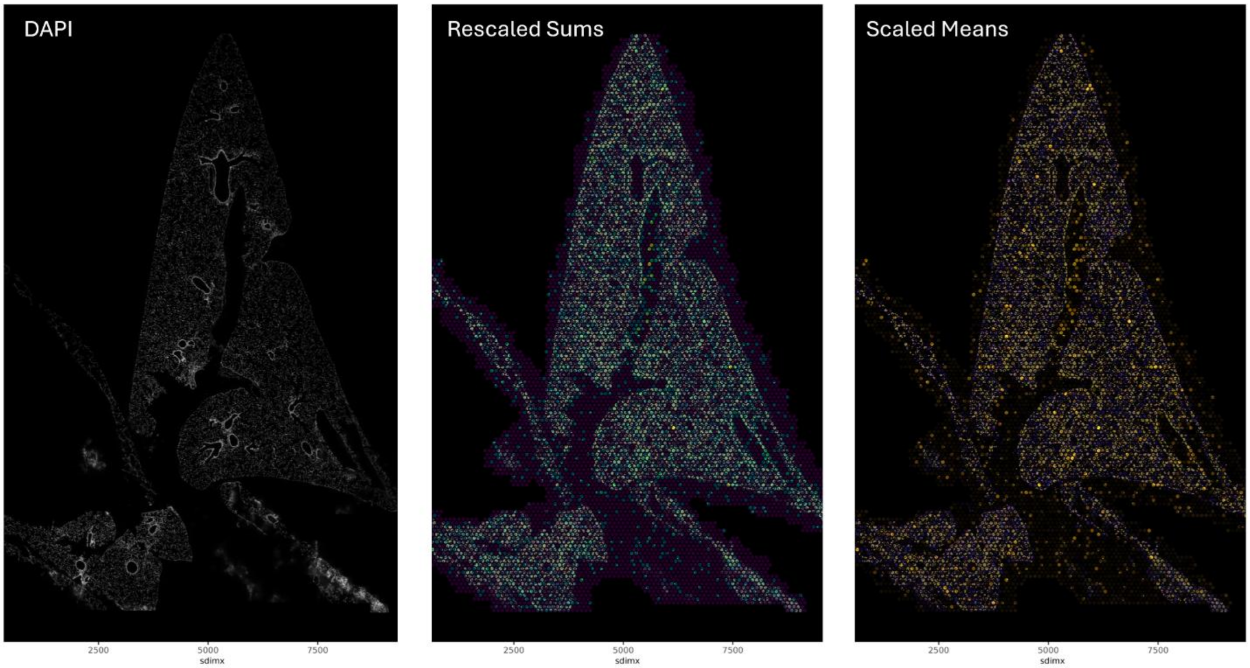


**Supplemental Figure S4.**

**Multiple implementations of hex-binning across samples.**

To locate potential megakaryocytes, a binning approach was used, whereby each image was subset into hexagonal compartments in an overlay (sample B pictured here). The color of each compartment was determined by enrichment of six MK marker genes, by looking at both rescaled sums, and scaled means (DAPI image on the far left for reference).
